# Targeting of MCL-1 in breast cancer associated fibroblasts reverses their myofibroblastic phenotype and pro-invasive properties

**DOI:** 10.1101/2021.09.24.461526

**Authors:** TL Bonneaud, L Nocquet, A Basseville, H Weber, M Campone, PP Juin, F Souazé

## Abstract

Cancer associated fibroblasts (CAF) are a major cellular component of epithelial tumors. In breast cancers in particular these stromal cells have numerous tumorigenic effects in part due to their acquisition of a myofibroblastic phenotype. Breast CAFs (bCAFS) typically express MCL-1. We show here that targeting this regulator of mitochondrial integrity using a specific BH-3 mimetic promotes fragmentation of these organelles without inducing cell death. MCL-1 antagonism in primary bCAFs directly derived from human samples mitigates myofibroblastic features and decreases expression of genes involved in actomyosin organization and contractility, associated with a cytoplasmic retention of the transcriptional regulator, Yes*-*Associated Protein (YAP). Such treatment decreases bCAFs ability to promote cancer cells invasion in 3D co-culture assays. These effects are counteracted by an inhibitor of the mitochondrial fission protein DRP-1, which interacts with MCL-1 upon BH3 mimetic treatment. Our findings underscore the usefulness of targeting MCL-1 in breast cancer ecosystems, not only to favor death of cancer cells but also to counteract the tumorigenic activation of fibroblasts with which they co-evolve.

The authors declare no conflict of interest.

## Introduction

In epithelial tumors, Cancer-associated fibroblasts (CAFs) are an important component of a hijacked stroma and may represent up to more than half of the whole breast tumor volume. These cells are composed of different heterogeneous and plastic populations with various supporting effect on cancer cells [1,2]. In breast cancer, different subsets of CAF have been described, differing in their expression markers, their secretions and their main characteristics (pro-tumor functions, immunoregulatory properties…) possibly linked to a different activation status. Subsets with myofibroblastic characteristics are enriched in tumor tissue and contribute to pathogenesis [3,4]. Myofibroblastic CAFs are indeed key mediators of fibrotic tumor stroma which is characterized by accumulation of extracellular matrix (ECM), increased tissue stiffness and breast cancer risk [5–7]. They have marked stress fibers and contract collagen, which underlies their pro-invasive properties [8,9]. The myofibroblastic characteristics of CAFs rely on the expression of a set of genes involved in the structure of the actomyosin cytoskeleton (among them the ACTA2 encoding for the myofibroblastic activation marker α-SMA) controlled by mechano-sensitive transcription factors, YAP-TEAD and MRTF-SRF [10,11]. These two pathways are mutually dependent and are closely regulated by the polymerization state of the actin cytoskeleton [11]. In addition to responding to mechanical stress, myofibroblast differentiation is also under the control of secreted factors [12].

Given the adverse effects of myofibroblastic CAFs on cancer progression and treatment resistance of cancer cells [13], their pharmacological manipulation and selective eradication within the tumor is therapeutically relevant [14–16]. Importantly, activated myofibroblasts are more refractory to cell death than other fibroblasts, and express higher levels of pro-survival proteins [17,18]. Fibroblasts from patients with scleroderma rely on BCL-XL or MCL1/BCL-XL for survival [17]. We ourselves established that MCL-1 mRNA expression positively correlates with stromal score in publicly available expression datasets from luminal breast cancers. CAFs derived from breast cancers (bCAFs), regardless of subtype, express MCL-1 at a higher level than in normal fibroblasts. This was in particular established using CAFs grown ex vivo in culture conditions that favor their myofibroblastic phenotype [13,19]. In these cells, MCL-1 exerts its canonical function as it prevents mitochondrial outer membrane permeabilization (MOMP) in conjunction with BCL-xL [13].

In this study, we explored whether the typical expression of MCL-1 in bCAFs endowed with a myofibroblastic phenotype may constitute a vulnerability. BH3 mimetics are small molecules developed to specifically antagonize the anti-apoptotic function of various BCL-2 homologs, including MCL-1 [20]. They bind to the BH3 binding grove of anti-apoptotic proteins, which is critical for their engagement of the BH3 domain of their pro-apoptotic, MOMP-promoting, counterparts. In numerous settings, inhibition of MCL-1 grove coincidentally stabilizes MCL-1, most likely because the turn-over of this labile protein is highly dependent upon its binding to endogenous proteins harboring a BH3, or BH3-like, domain [21–23]. BH3 mimetics were initially conceived to pharmacologically favor MOMP and apoptosis in cancer cells, which are characterized by frequent alterations in the expression and activity of BCL-2 homologs. MCL-1 in particular is frequently overexpressed in many cancers, including breast cancers, and this was established to contribute to resistance to the cytotoxic effects of chemotherapy, radiotherapy and BH-3 mimetics selectively targeting other BCL-2 homologs [13,20,24,25]. We also established that in luminal breast cancers MCL-1 expression in tumor cells is extrinsically favored by paracrine effects of bCAFs. This defines MCL-1 as a target in stroma-influenced breast cancers, and advocates for a comprehensive investigation of the effects of its targeting in distinct cellular components, such as myofibroblastic CAFs, of these tumor ecosystems. We describe here that targeting MCL-1 with a BH-3 mimetic fails to induce efficient apoptosis in bCAFs. However, treatment leads to a loss of myofibroblastic features of these cells, which we ascribe to a role of MCL-1-dependent mitochondrial and actomyosin dynamics in the bCAFs tumorigenic phenotype.

## Material and methods

### Cell culture and reagents

Fresh human mammary samples were obtained from treatment naive patients with invasive carcinoma after surgical resection at the Institut de Cancérologie de l’Ouest, Nantes/Angers, France. As required by the French Committee for the Protection of Human Subjects, informed consent was obtained from enrolled patients and protocol was approved by Ministère de la Recherche (agreement n°: DC-2012-1598) and by local ethic committee (agreement n°: CB 2012/06). Breast cancer associated fibroblasts (bCAFs) isolation and characterization were previously described in Louault et al, 2019 [13].

For the CRISPR Cas9-induced knock-out (KO) primary bCAFs, single guide (sg) RNA sequences targeting human genes were designed using the CRISPR design tool (http://crispor.tefor.net). The guide sequences CGCGGTGACGTCGGGGACCT were cloned in the plentiCRISPRV2 vector that was a gift from Feng Zhang (Addgene plasmid # 52961)[26]. Cells were selected using 1 μg/ml puromycin and protein extinction were confirmed by immunoblot analysis.

The human breast cancer cell lines T47D were purchased from American Type Culture Collection (Bethesda, MD, USA), GFP was introduced by lentiviral infection with PFG12 (insert EGFP) from Addgene. FG12 was a gift from David Baltimore (Addgene plasmid # 14884; http://n2t.net/addgene:14884; RRID:Addgene_14884).

BH3 mimetics, S63845 and ABT-199 (Chemieteck, Indianapolis, IN, USA), AZD-5991 and A1331852 (MedChemExpress, NJ, USA), ABT-737 (Selleckchem, Houston, TX, USA) and Q-VD-OPh (Sigma-Aldrich, Saint-Louis, MO, USA) and Mdivi-1 (Sigma-Aldrich, Saint-Louis, MO, USA) were dissolved in DMSO and used at the indicated concentrations.

### Apoptosis assays

The activity of caspase-3/7 was measured by Caspase-Glo-3/7 assay kit according to the manufacturer’s instructions (Promega). Cell death was assessed by an Annexin-V FITC binding assay (Miltenyi, France) performed according to manufacturer’s instructions. Flow-cytometry analysis was performed on an Accuri C6 flow cytometer (BD biosciences San Jose, CA, USA).

### Immunocytochemistry

Cells were fixed in PBS containing 4% paraformaldehyde/4% sucrose for 15 min. Cells were permeabilized for 5 min at room temperature in 0.25% Triton-X-100 in PBS, washed twice with PBS, and incubated for 30 min at 37°C in PBS containing 10% BSA. Cells were incubated overnight at 4°C with primary antibodies diluted in PBS containing 3% BSA. Antibodies used were as follows: mouse anti alpha-SMA (1:200, Invitrogen, Carlsbad, CA, USA); Myosin-IIB (1:1000, BioLegend, San Diego, CA, USA); YAP1 (1:50, Proteintech, Manchester, UK); TOM20 (1:80, Santa Cruz Biotech., Dallas, TX, USA); Mcl-1 (1:800, Cell Signaling Technology, Danvers, MA, USA); Vimentin (1:100, Merck, Darmstadt, Germany). Actin labeling were performed with Alexa 488-conjugated Phalloidin (1:100, Cell Signaling Technology, Danvers, MA, USA). After washing, cells were incubated for 90 min at room temperature with the appropriate Alexa 488-conjugated secondary antibodies diluted in PBS containing 3% BSA. Cells were washed with PBS and mounted with ProLong Diamond Antifade Reagent with DAPI (Invitrogen, Carlsbad, CA, USA). Fluorescence images were acquired with Nikon A1 Rsi Inverted Confocal Microscope (Nikon, Tokyo, Japan) with NIS-Elements software (Nikon). Images set position were generated randomly through NIS-Elements JOBS module (Nikon).

### Immunoblot analysis

Cells were resuspended in lysis buffer (1% SDS; 10 mM EDTA; 50 mM Tris-Hcl pH 8.1; 1 mM PMSF; 10 μg/ml^−1^ aprotinin; 10 μg/ml^−1^ leupeptin; 10 μg/ml^−1^ pepstatin; 1 mM Na3VO4 and 50 mM NaF). For western blotting, following SDS–PAGE, proteins were transferred to 0.45 μM nitrocellulose membranes using Trans-Blot® Turbo™ Transfer System Cell system (Bio-Rad). The membrane was then blocked in 5% nonfat dry milk TBS 0.05% Tween 20 and incubated with primary antibody overnight at 4°C. Blots were incubated with the appropriate secondary antibodies for 1 h at room temperature and visualized using the Chemi-Doc XRS+ system (Bio-Rad). Primary antibodies used were anti-BCL-X_L_ (abcam) (ab32370), anti-MCL-1 (Santa Cruz) (sc-819), anti-BCL-2 (Dako) (M0887), anti-Bax (Dako) (A3533), anti-Bak (Cell signaling) (3814), anti-Tom20 (ab186734) (Abcam) anti-β-ACTIN (Millipore) (MAB1501R).

### Co-Immunoprecipitation

CAFs were collected in lysis buffer (CHAPS 1%, TRIS-HCl 20mM pH=7.5, NaCl 150mM, EDTA 1mM pH=8.0, proteases and phosphatases inhibitors). Co-Immunoprecipitation was performed using PureProteome™ Protein A Magnetic Beads (Millipore, LSKMAGA10). Protein extracts were first precleared using 10μl of beads for 200μg of proteins (1-hour incubation at 4°C). 2 μl of anti-MCL-1 antibody (Cell signaling mAb #94296) and 10μl of beads were then used for 200 μg of proteins following manufacturer’s instructions (direct protocol). For western blotting, DRP-1 antibody (MA5-26255, Thermo Fisher Scientific, Waltham, MA, USA) was used as primary antibody (1:1000) with the appropriate secondary antibodies and the same anti-MCL-1 primary antibody (1:1000) and Clean-Blot™ IP Detection Reagent (HRP) (Thermo Scientific #21230) was used as secondary antibody (1:1000).

### seahorse

OCR measurements were performed using the Seahorse XF HS Mini Analyzer (Seahorse Bioscience). Cells were plated in duplicate in 8-well seahorse plate (7 000 cells per well) and treated with S63845 500nM for 18h. Then, medium was replaced by glutamine/glucose/pyruvate-free DMEM (no sodium bicarbonate) adjusted to pH∼7.4 and incubated for 45 min at 37% in a CO_2_-free incubator. OCR was normalized to cell number ratio determined by cell counting at the end of the experiment.

### Analysis of mRNA expression data in CAFs

3’seq-RNA Profiling protocol is performed according to Soumillon et al [27]. The libraries are prepared from 10 ng of total RNA in 4μl. The mRNA poly(A) tails are tagged with universal adapters, well-specific barcodes and unique molecular identifiers (UMIs) during template-switching reverse transcription. Barcoded cDNAs from multiple samples are then pooled, amplified and tagmented using a transposon-fragmentation approach which enriches for 3′ends of cDNA: 100ng of full-lentgth cDNAs are used as input to the Nextera DNA Sample Prep kit (ref FC-121-1030, Illumina) which enriches for 3′ends of cDNA. Size library is controlled on 2200 Tape Station Sytem (Agilent Technologies). A library of 350– 800 bp length is run on an Illumina HiSeq 2500 using a Hiseq Rapid SBS Kit v2-50 cycles (ref FC-402-4022) and a Hiseq Rapid PE Cluster Kit v2 (ref PE-402-4002) according to manufacturer’s protocol (Denaturing and Diluting Libraries for the HiSeq® and GAIIx, Part # 15050107 v03 protocol, Illumina).

Raw fastq pairs match the following criteria: the 16 bases of the first read correspond to 6 bases for a designed sample-specific barcode and 10 bases for a unique molecular identifier (UMI). The second read (58 bases) corresponds to the captured poly(A) RNAs sequence. We perform demultiplexing of these fastq pairs in order to generate one single-end fastq for each of the 96 samples. These fastq files are then aligned with bwa to the reference mRNA refseq sequences and the mitochondrial genomic sequence, both available from the UCSC download site.

Gene expression profiles are generated by parsing the alignment files (.bam) and counting for each sample the number of UMIs associated with each gene. Reads aligned on multiple genes, containing more than three mismatches with the reference sequence or having a polyA pattern are discarded. Finally, a matrix containing the counts of all genes on all samples is produced. The expression values, corresponding to the absolute abundance of mRNAs in all samples, is then ready for further gene expression analysis. The R package deseq2 [28] is then used for the differential analysis. Batch effect was corrected according to CAF patient origin using Limma package. Differentially expressed genes were calculated using ebayes function (Limma) and filtered in with an adjusted p-value inferior to 0.05 and a log2 fold-change superior to 1. Heatmap was built with complexHeatmap package using euclidean distance and ward D2 clustering method. Pathway enrichment analysis was done using clusterprofiler package.

### Collagen gel contraction assay

Fibroblasts (100 000 cells) were re-suspended in 500 μl collagen type I suspension (1 mg/ml) (BD/corning) (354249) in DMEM supplemented with 5% FBS. Cell suspension was cast into each well of 24-well tissue culture plate and incubated at 37°C for 1 h in order to facilitate gelation. Following this, gels were released from the surface of the culture well using a sterile tip. Contraction between the different conditions was evaluated at 3 h.

### RNA isolation and quantitative real-time PCR

Total RNA was isolated using Nucleospin RNA (Macherey-nagel, Hoerdt, France) and transcribed into cDNA by Maxima First Strand cDNA synthesis Kit (Thermo scientific). Quantitative RT-PCR (qPCR) was performed using the EurobioGreen qPCR Mix Lo-Rox with qTOWER (Analityk-jena, jena, Germany). Reaction was done in 10 μl final with 4 ng RNA equivalent of cDNA and 150 nM primers. Relative quantity of mRNA was estimated by Pfaffl method (Pfaffl *et al*., Nucleic Acids Res, 2001) and normalized on the average relative quantity of two housekeeping genes.

RPLP0 5’-AACCCAGCTCTGGAGAAACT/CCCCTGGAGATTTTAGTGGT-3’

GAPDH 5’-CAAAAGGGTCATCATCTCTGC/AGTTGTCATGGATGACCTTGG-3’

ACTA2 5’-CTATGCCTCTGGACGCACAACT/CAGATCCAGACGCATGATGGCA-3’

MYH2 5’-GTCTGCCAACTTCCAGAAGC/CAGCTTGTTCAAATTCTCTCTGAA-3’

MYH10 5’-GACTGAGGCGCTGGATCTGT/AAAGCAATTGCCTCTTCAGCC-3’

MYLK 5’-GGTGACATGGCACAGAAACG/AGCTGCTTCGCAAAACTTCC-3’

COL11A 5’-TGGTGATCAGAATCAGAAGTTCG/AGGAGAGTTGAGAATTGGGAATC-3’

ITGA11 5’-CGGCCTCCAGTATTTTGGCT/GGAGGCTGGCATTGATCTGA-3’

LBH 5’-GCCCCGACTATCTGAGATCG/GCGGTCAAAATCTGACGGGT-3’

ELN 5’-GTCCTCCTGCTCCTGCTGT/CTCCTCCTCCAAGGGCTC-3’

### 3D spheroid Collagen invasion

3D-spheroids were obtained by re-suspending cells at a concentration of 1.25 10^4^ cells/ml in 20% Methyl-cellulose/80% DMEM solution supplemented with 10% FBS. Then, 100μL per well were distributed in 96-well-conical-plates (non-treated surface) and centrifuged 1min at 200g. Spheroids were harvested the next day and transferred in a collagen type I suspension (2 mg/ml) (BD/corning) (354249) supplemented with 1% FBS in presence of the indicated treatments. After collagen polymerization, DMEM 1% FBS (+/-treatment) were added to the top of each well to prevent any gel desiccation. Invasion was monitored by fully automated *Dmi6000b Wide Field* Fluorescence *Microscope* (Leica Microsystems, Wetzlar, Germany) during 48h.

## Results

### MCL-1 targeting in bCAFs promotes DRP-1 dependent mitochondrial fragmentation without significantly increasing cell death rates

We showed that high expression level of MCL-1 was a hallmark of activated bCAFs (Louault et al, 2019). bCAFs systematically express the anti-apoptotic protein MCL-1 at a high level, while expression of BCL-xL is sporadic and that of BCL-2 barely detectable. To investigate whether this set of expressions corresponds to specific survival dependencies, we performed pharmacological BH-3 profiling of primary cultures of bCAFs (derived from human breast cancer samples). We treated these cells with combinations of BH-3 mimetics targeting MCL-1 (S63845), BCL-2 (ABT-199), BCL-xL (A1331852) or BCL-2/BCL-xL/BCL-w (ABT-737) and measured cell death rates (using Annexin V binding as a readout). This revealed that bCAFs survival does not rely on the function of a single anti-apoptotic protein (and a fortiori on that of MCL-1 alone) but rather on the combined functions of MCL-1 and BCL-xL (Figure 1A). Acordingly, in contrast to combined targeting (using for instance S63845 + ABT-737) which triggered robust caspase 3/7 activation, targeting MCL-1 only resulted in not significant caspase-3/7 activation (Figure1B, S63845).

**Figure 1.**
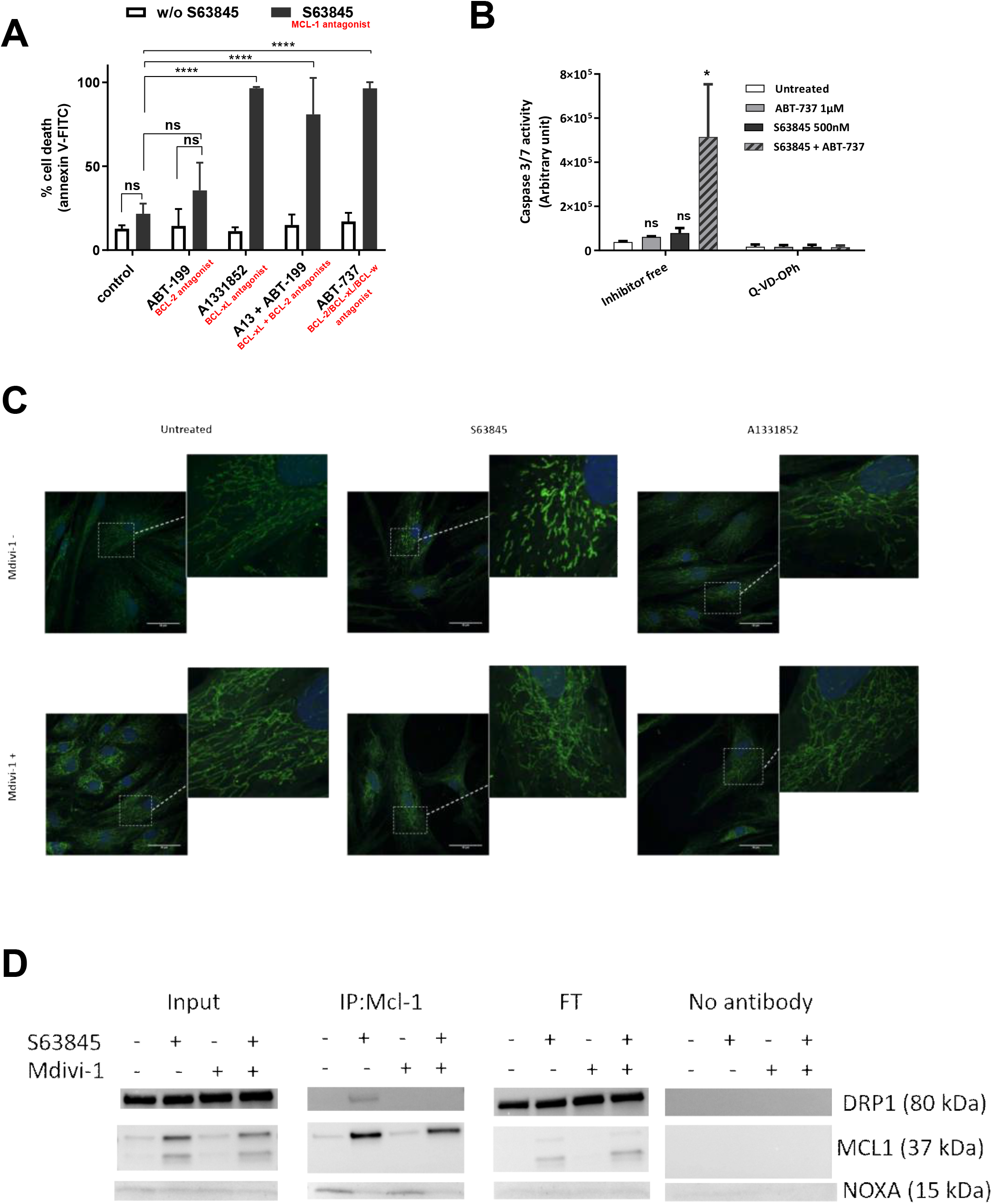
Targeting MCL-1 in bCAFs induces DRP-1 dependent mitochondrial fragmentation without significant cell death. **A** Three different primary cultures of CAFs were treated with ABT-199 (1μM), A1331852 (100nM), ABT-199 (1μM) +A1331852 (100nM) or ABT-737 (1μM) with or without S63845 (500nM) for 72 hours in DMEM containing 1% FBS, apoptosis was measured by Annexin-V flow cytometry. Data are means ± SEM from three independent experiments. Two-way ANOVA, ****P<0.0001, ns: not significant. **B** Caspase 3/7 activity was measured by caspase Glo assay and expressed in arbitrary unit. Data are means ± SEM from three independent experiments. Two-way ANOVA. *P<0.05, ns: not significant **C** Mitochondrial fission was detected by TOM20 staining in CAFs treated or not with S63845 500nM or A1331852 100nM for 18 hours in presence or not of Mdivi-1 50μM. Scale bar = 50 μm. **D** Immunoprecipitation of Mcl-1 and Western blotting for Mcl-1, Drp1 or Nox-A in CAFs treated or not with S63845 in presence or not of Mdivi-1.

In addition to MOMP, MCL-1 was shown to regulate the dynamics between mitochondrial fusion and fission in human pluripotent stem cells (hPSCs) and in hPSCs-derived cardiomyocytes [29], and S63845 treatment of cardiomyocytes was shown to promote mitochondrial fission [30]. A more thorough investigation of the impact of S63845 treatment on the mitochondrial network of bCAFs showed that it induced mitochondrial fragmentation in bCAFs whereas BCL-xL antagonist (with A1331852) did not (Figure 1C). This mitochondrial ultrastructural defect did not coincide with detectable changes in oxidative phosphorylation (measured with a Seahorse, Supplementary Figure 1A) and mitochondrial mass (evaluated by total TOM-20 expression Supplementary Figure 1B). The effect of S63845 was prevented by co-treatment with Mdivi-1, an inhibitor of dynamin-related protein 1 (DRP-1), indicating that it relies on activation of the endogenous mitochondrial fission machinery of which this protein is a critical member (Figure 1C). Mechanistically, and in line with previous reports in numerous other cell types, bCAFs treatment with S63845, but not with A1331852, enhanced MCL-1 protein levels (Supplementary Figure 1C and D). MCL-1 accumulated preferentially at the mitochondria as illustrated by the colocalization of MCL-1 and TOM20 seen in immunocytochemistry (Supplementary Figure 1D). Based on studies conducted in hPSCs and hPSC-derived cardiomyocytes, we evaluated interactions between MCL-1 and DRP-1 by co-immunoprecipitation experiments. We show detectable amounts of DRP-1 co-immunoprecipitated with MCL-1 following treatment with S63845 (Figure 1D), note that S63845 treatment prevented MCL-1/NOXA interaction. Co-treatment with Mdivi-1 did not prevent accumulation of MCL-1 protein levels but prevented co-immunoprecipitation with DRP-1 (Figure 1D). Taken together, these data indicate that treatment with MCL-1 specific BH3-mimetic increases DRP-1 dependent mitochondrial fission in CAFs, which coincides with enhanced interactions between endogenous Mcl-1 and Drp1.

### MCL-1 targeting influences transcriptional programs involved in actomyosin organization/contractility and mitigates myofibroblastic features

To further document the biological effects of MCL-1 targeting on bCAFs, we performed a differential gene expression (DGE) analysis on RNA-seq data from six different primary cultures of bCAF (derived from 3 TNBC and 3 Luminal breast cancers) treated or not with S63845 (500nM, 18h). As observed in a volcano plot representation (Figure 2A), we identified 23 genes that were significantly upregulated and 20 genes that were significantly downregulated after S63845 across all bCAFs. Importantly, changes in the expressions of these 43 genes induced by S63845 prevail over inter-sample variations. Indeed, hierarchical clustering of untreated and treated bCAFs according to these 43 differentially expressed genes (See Heatmap in Figure 2B) split them into two clusters defined by the treatment received. No additional clustering was observed according to cancer subtype origin (TNBC or Luminal). We confirmed by RT-PCR the S63845-induced down-regulation of a selected set of genes (MYH10, MYH2, ACTA2, COL11A1, ITGA11, LBH and MYLK) and showed that this effect is reversed by Mdivi-1 for most tested genes (except ITGA11 and MYLK) (Figure 2C).

**Figure 2.**
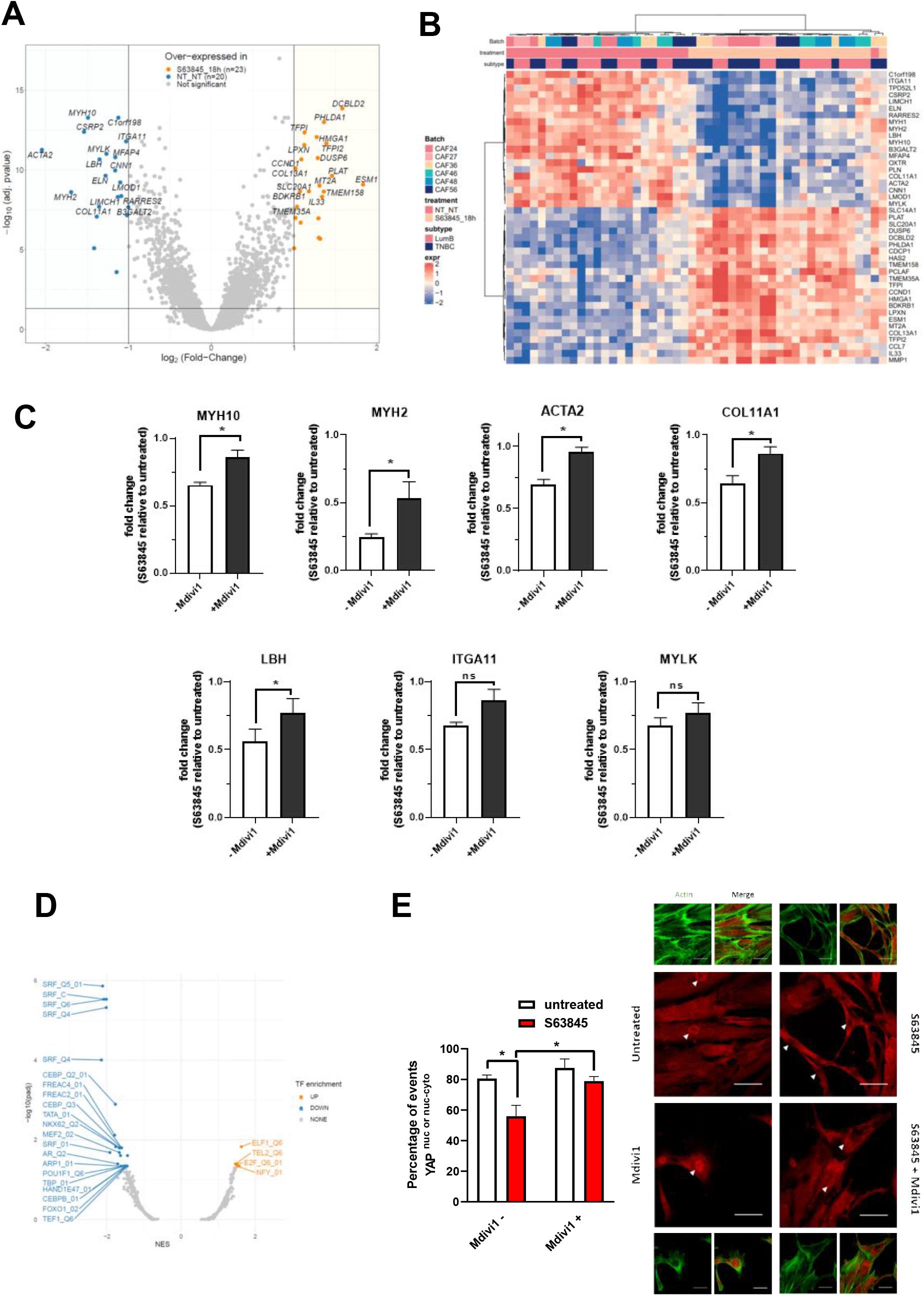
MCL-1 targeting influences transcriptional programs involved in actomyosin organization/contractility. **A** Volcano plot displaying differential expressed genes between S63845 treated and non-treated primary culture of CAFs from luminal B (n=3) and triple negative (n=3) breast cancers. The vertical axis (y-axis) corresponds to the mean expression value of log 10 (q-value), and the horizontal axis (x-axis) displays the log2 fold change value. The red dots represent the up-regulated expressed transcripts; the blue dots represent the transcripts whose expression is downregulated. **B** A hierarchically clustered heatmap showing the expression patterns of the 23 most extreme DE mRNA. Red and blue represent up- and down-regulated expression in CAFs treated or not with S63845. Color density indicating levels of fold change was displayed. **C** qRT-PCR of MYH10, MYH2, ACTA2, COL11A1, ITGA11, LBH and MYLK mRNA in CAFs treated or not by S63845 in presence or not of Mdivi-1. Mean and SEM of four or five independent experiments are represented as fold change of mRNA level in CAFs treated with S63845 related to untreated. Student t-test, * P<0.1 **D** Volcano plot of the enriched transcription factor. TF: transcription factor. NES, normalized enrichment score. **E** Quantification (percentage of cells positive for nuclear YAP or nuclear and cytosolic YAP) (left panel) and confocal image (right panel) of YAP staining (red) and FITC-phalloidin (green) in CAFs treated or not with S63845 in presence or not of Mdivi-1. Randomly generated images set names were randomized for analysis. Around sixty to one hundred cells were analyzed per condition. Counting events were done manually through NIS-Elements software (Nikon software). Fluorescence intensity and positivity staining were determined by positive and negative control comparison, dependent on the experiment. The nuclei were counterstained with DAPI (blue). Scale bar = 50 μm. Two-way ANOVA, *P<0.05.

Transcription factor enrichment analysis of S63845-induced differential gene expression hinted on SRF as a candidate transcription factor possibly involved in the observed down regulations (Figure 2D). However, we could not detect any change in the nuclear localization of SRF or of its cofactor, MRTF, upon treatment (data not shown). SRF/MRTF-regulated genes were shown to co-depend on YAP in bCAFs [11]. We found that the proportion of bCAFS exhibiting nuclear localization of YAP (including nuclear and cytosolic) was decreased upon treatment and reversed in the presence of Mdivi1 (Figure 2E). Without ruling out the involvement of additional transcriptional factors, this argues that cytoplasmic retention of YAP may contribute, at least in part, to the transcriptional effects of S63845.

Gene Ontology (GO) pathway enrichment analysis based on DGE results indicated that MCL-1 targeting in bCAFs was mainly associated with a decreased expression of genes involved in extracellular matrix and actomyosin structure organizations and in contractility (Figure 3A). Among these genes, the majority is known to be involved in myofibroblastic differentiation and are *bona fide* CAF markers such as ACTA2 (coding for α-SMA), COL11A1 or ITGA11 CAF markers (Figure 2E and 3A-C). To validate that MCL-1 targeting influences bCAFs myofibroblastic phenotype we evaluated α-SMA protein expression, contractile force and surface area upon treatment with S63845. This treatment induced a decrease in α-SMA expression (Figure 3B), inhibited collagen contraction ability (Figure 3C) and reduced CAFs surface area (Figure 3D).

**Figure 3.**
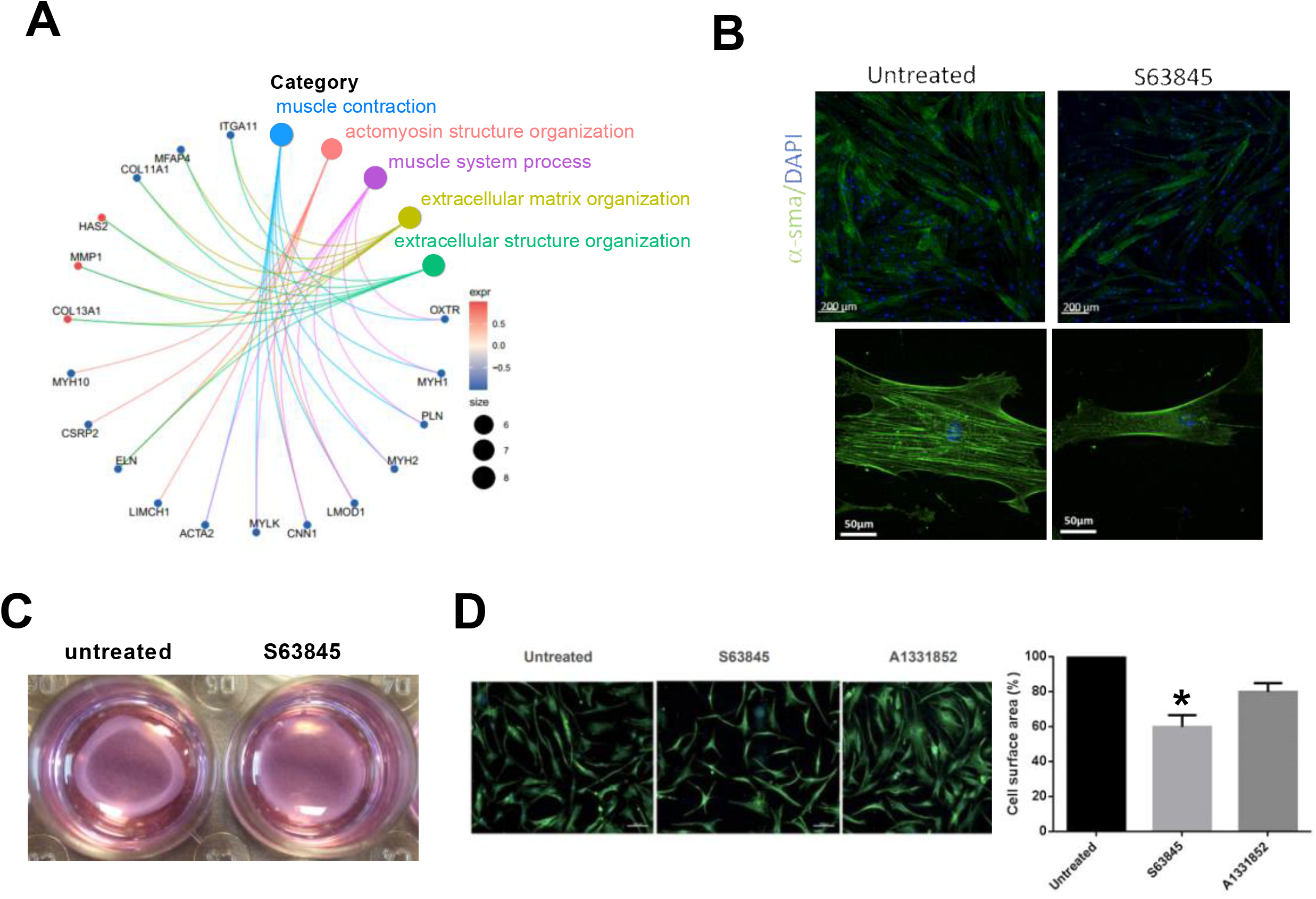
MCL-1 antagonist mitigates myofibroblastic features. **A** Gene Ontology (GO) pathway enrichment analysis based on DGE results. **B** Immunofluorescence of -smooth muscle actin (α-SMA) (green) in CAF treated or not with S63845 for 18 hours, nuclei were stained in blue (4’,6-diamidino-2-phenylindole, DAPI). **C** Images of CAFs contraction of collagen gels. **D** Cells surface area of CAFs determined by Vimentin immunostaining. Cells area were measured with imageJ software, results are expressed relative to untreated cells. Student t-test, * P<0.1.

### MCL1 targeting induces changes in actomyosin structure organization and loss of myofibroblastic features of bCAFs in a DRP-1 dependent manner

The effects of MCL-1 targeting on MYH10, MYH2, ACTA2, COL11A1 and ITGA11 mRNA down-regulation (Figure 2E) and on YAP cytoplasmic retention (2D) were reversed by Mdivi-1 treatment. To further document a link between MCL-1 regulated mitochondrial dynamics and myofibroblastic features, we co-evaluated actin stress fibers (as markers of myofibroblast features, stained by F-actin marker, Phaloidin) and mitochondrial networks (distinguished between fragmented and fused by immunolabelling of the outer mitochondrial membrane protein TOM20-Figure 4) in bCAFS cultured in the absence or presence of S63845. Around 80% of untreated bCAF populations showed fused mitochondria associated with marked stress fibers. Half of the remaining 20% exhibiting fragmented mitochondria were negative for stress fibers (Figure 4A, untreated). Following treatment with S63845, a shift towards a major proportion of CAFs (60-80%) with fragmented mitochondria (of which 40-60% were negative for stress fibers) occurred (Figure 4A). This was recapitulated by treatment with another MCL-1 targeting BH3 mimetic (AZD5991) but not by a BCL-xL targeting BH3 mimetic (A1331852). The effects of MCL-1 selective BH3 mimetics were counteracted by Mdivi1.

**Figure 4.**
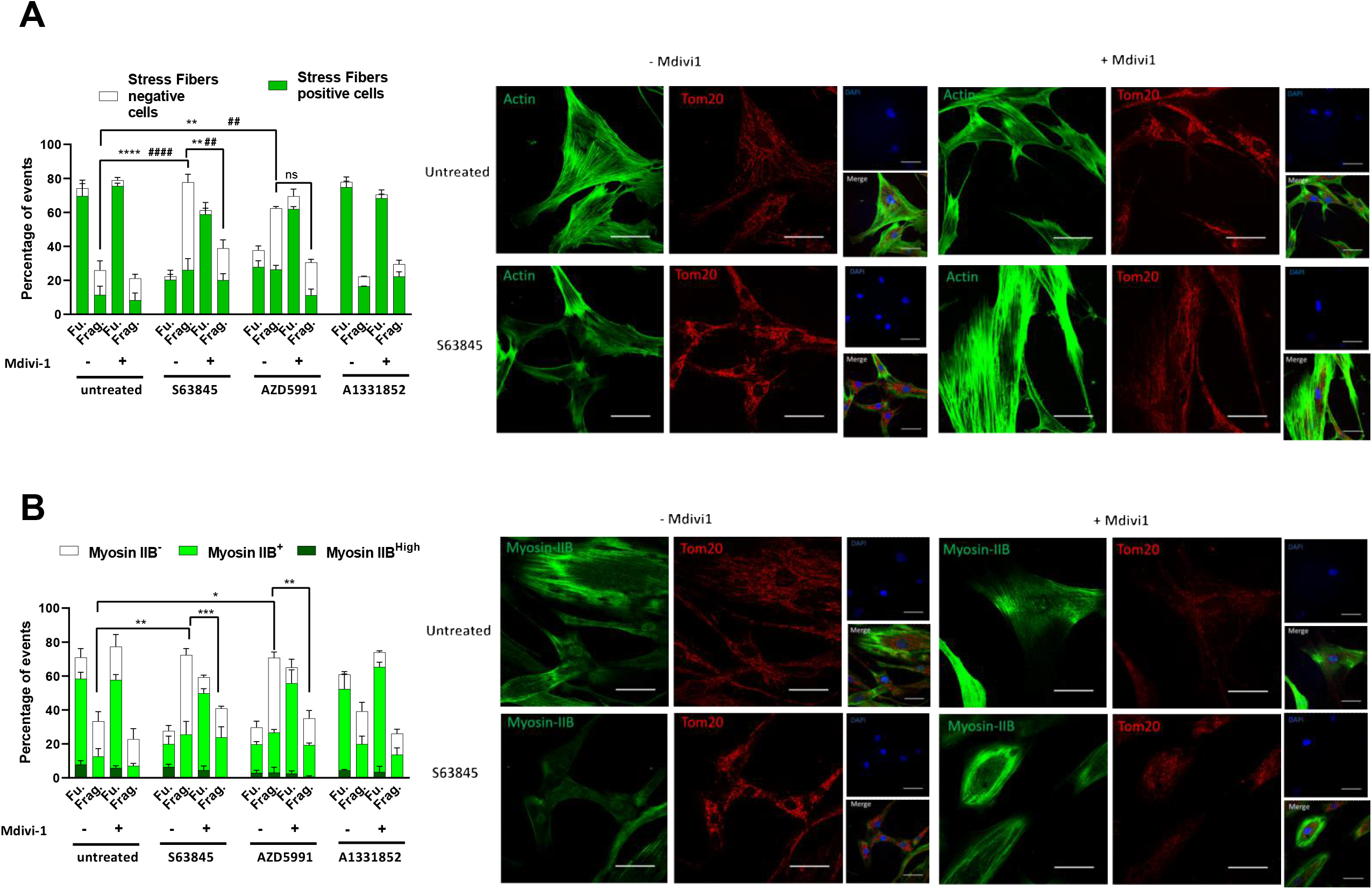
MCL-1 targeting induces changes in actomyosin structure organization in a DRP-1 dependent manner. **A-B** Quantification (left panel) and confocal image (right panel) of (A) F-actin staining with FITC-phalloidin (green) or (B) Myosin IIB staining (green) and immunofluorescence of TOM-20 (red) in CAFs treated or not with different BH3 mimetics (as indicated) in presence or not of Mdivi-1. Randomly generated images set names were randomized for analysis. Around sixty to one hundred cells were analyzed per condition. Counting events were done manually through NIS-Elements software (Nikon software). Fluorescence intensity and positivity staining were determined by positive and negative control comparison, dependent on the experiment. The nuclei were counterstained with DAPI (blue). Scale bar = 50 μm, (A) Two-way ANOVA, ^####^ P<0.0001, ^##^P<0.01 for stress fibers negative cells, **** P<0.0001 ** P<0.01 for fragmented mitochondria, ns: not significant. (B) **** P<0.0001, ** P<0.01, * P<0.1

We also measured the effect of MCL-1 targeting compounds on myosin-IIB (MYH10) expression since this protein has an established role in stress fibers organization [31] and since its mRNA expression was found downregulated by S63845 (Figure 2A-C). S63845 (or AZD5991)-induced mitochondrial fragmentation was accompanied by a decrease in Myosin-IIB expression in bCAFs with 60% of CAFs not expressing it (mostly exhibiting fragmented mitochondria) versus 30% in the control (Figure 4B). Mdivi-1 co-treatment prevented these effects (Figure 4B). A1331852 had no effect on myosin-IIB expression (Figure 4B).

These results put forth a link between mitochondrial dynamics and organization of the actomyosin cytoskeleton of bCAfs, bringing further support to the notion MCL-1 targeting contributes to a loss of the myofibroblastic phenotype of bCAFs by promoting DRP-1 dependent mitochondrial fragmentation.

### MCL-1 targeting in bCAFs decreases invasiveness and pro-invasive effects on cancer cells

We finally investigated the consequence of S63845-induced phenotypic changes on the invasive properties of bCAFs. To this end, we analyzed invasion of bCAFs spheroid treated or not with S63845 in type I collagen matrix. As shown in figure 5A, whereas untreated bCAFs rapidly invaded the collagen matrix in these 3D assays, MCL-1 targeting significantly reduced bCAFs invasion. As bCAFs were described to promote cancer cells invasion [8], we also investigated fibroblast-led invasion of breast cancer cells and the influence S63845 treatment in 3D coculture assays. Cancer cells were engineered to express GFP as a discriminating marker and we chose to use luminal T47D breast cancer cells that, under our experimental conditions, did not invade the collagen matrix when grown as spheroids in the absence of bCAFs (Figure 5B, Top). Upon coculture with bCAFs as 3D-multicellular tumor spheroids in type I collagen, GFP-positive T47D cancer cells invaded collagen and this was blocked by S63845 treatment (Figure 5B, Middle and Bottom). Loss of myofibroblastic features and of contractility noted upon S63845 treatment are accompanied by a decrease in the pro-invasive properties of bCAFs.

**Figure 5.**
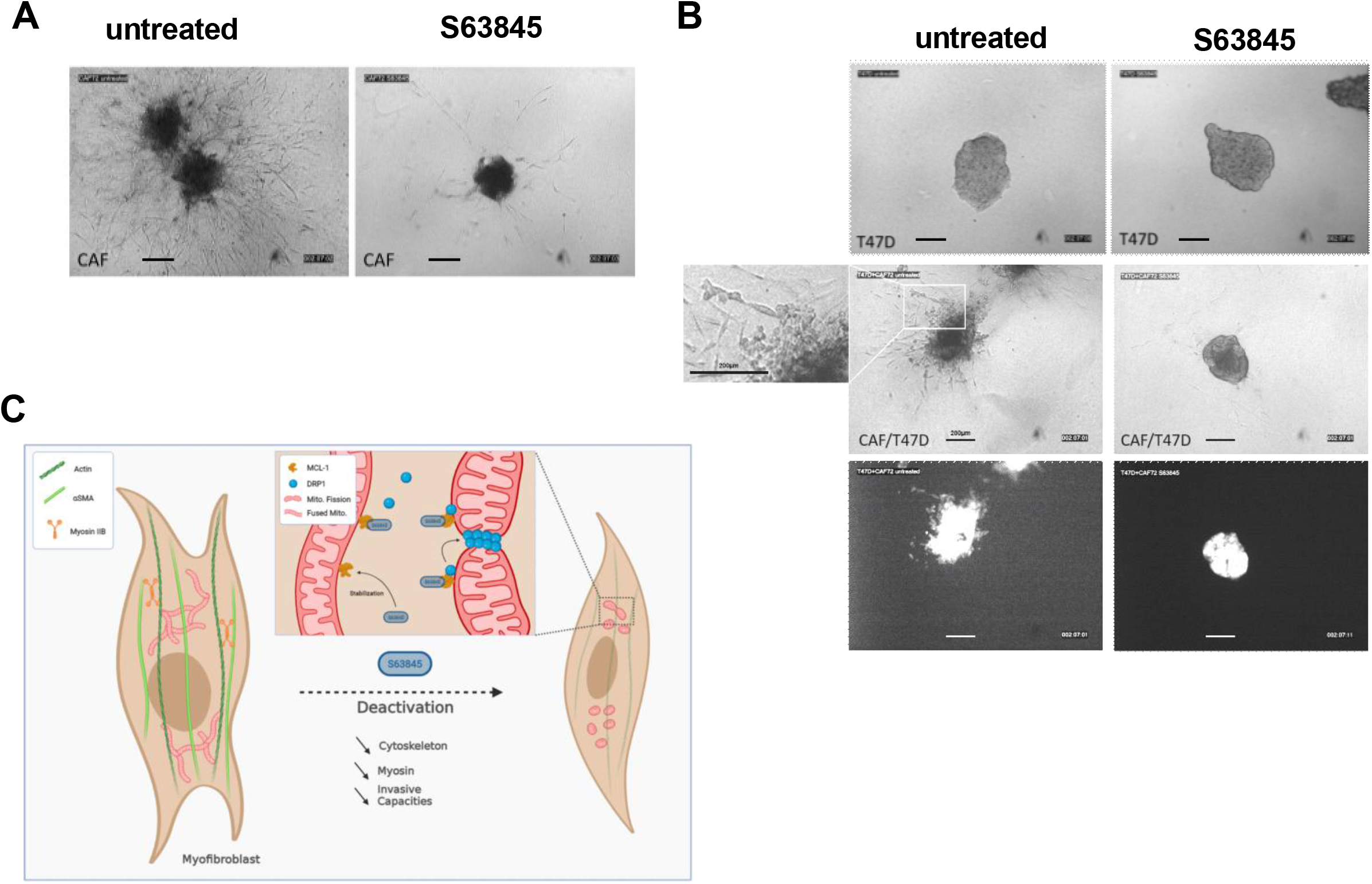
MCL-1 targeting in bCAFs decreases invasiveness and pro-invasive effects on cancer cells. **A** CAFs treated or not with S63845 **B** T47D cancer cells (top) or multicellular CAF + T47D-eGFP (middle and bottom) 3D spheroid invasion into collagen in presence or not of S63845 (500nM) was monitored up to 48h using microscopy time-lapse. Representative images of tumor cell invasion are shown. **C** Schematic model representing the influence of S63845 on the myofibroblastic phenotype of CAFs (Created with BioRender.com).

## Discussion

We herein present a novel and unexpected effect of a MCL-1 antagonist on cancer-associated fibroblasts, which reverses their myofibroblastic phenotype and decreases their pro-invasive capacity. Numerous studies showed that tumor intrinsic deregulation of MCL-1 expression/activity promotes breast cancer progression and treatment resistance. We also previously showed that in luminal breast cancers, MCL-1 expression in tumor cells was extrinsically favored by neighboring CAFs, which themselves exhibit high MCL-1 expression [13]. Our study thus brings support to the notion that a MCL-1 antagonist may improve the treatment of breast cancer [20,25] while putting forth an effect not only on cancer cells but also on non-malignant cells that interact with them. One implication is that the therapeutic potential of such a BH3 mimetic may be preclinically overlooked if reductive models using only tumor cells are used. Our study highlights instead that diverse cellular components of the microenvironment may experience an intense pressure on BCL-2 family dependent mechanisms regulating mitochondrial integrity/organization during tumor growth. This advocates for the use of assays that include cellular components of the microenvironment to investigate the effects of targeted agents such as BH3 mimetics, but also of cytotoxic compounds that indirectly impact on the BCL-2 network.

While in some models BH3 mimetics antagonizing MCL-1 efficiently counter tumor progression [32], the sole targeting of MCL-1 in solid tumors proved little efficiency to trigger cell death on its own, both in xenograft and PDX immunodeficient mice models [25,33]. Likewise, we observed that treatment of bCAFs with S63845 only induced moderate apoptotic rates. Moreover, treatment with a BCL-xL antagonist (A13), which alone did not induce bCAF death, did so in combination with S63845. Inefficient killing of the latter as a single agent may thus ensue from dynamic sequestration by BCL-xL of proapoptotic proteins released from MCL-1 upon treatment. We cannot rule out, however, that S63845 inefficiency when used alone may stem from the lack of binding of pro-apoptotic proteins to MCL-1 at baseline, (leaving “empty” MCL-1 available to engage pro-apoptotic proteins released from BCL-XL by A13, and to protect from cell death). In all cases, our functional studies show that MCL-1 combines its anti-apoptotic activity with that of BCl-xL to maintain bCAFs survival. Consistently, CRISPR-Cas9 engineered knock down of MCL-1 in bCAFs dramatically sensitized them to a BCL-xL inhibitor (Supplementary Figure 2A).

Even though the pro-apoptotic effects of MCL-1 targeting are restrained by BCL-xL activity, MCL-1 targeting by itself modifies the tumorogenic phenotype of bCAFs by mitigating their myofibroblastic features and invasive properties. We ascribed this process to induction of mitochondrial fragmentation, as changes induced in the actomyosin network by MCL-1 targeting where counteracted by inhibition of DRP-1. Our study evokes reports indicating that MCL-1 is not only an anti-apoptotic protein determining life death decisions but also a more subtle regulator of mitochondrial dynamics and of mechanical properties in various cell types. A role for MCL-1 in the maintenance of mitochondrial structure and of cellular differentiation was described in cardiomyocytes, and stem cells [29,30]. Moreover S63845 was already described to promote mitochondrial fission and to prevent cardiomyocyte function and contractility [30]. This latter effect, which corroborates our study, may be the basis for the dose limiting cardiotoxic effect of MCL-1 antagonists when used in the clinic [34]. The therapeutic window for exploiting the effects of MCL-1 targeting on a myofibroblastic tumor stroma described here may thus be narrow, unless specific tumor delivery strategies intervene.

Our data argue that the molecular basis for the effects of S63845 in bCAFs are not canonical, in that they do not seem to rely on the release of one or more pro-apoptotic proteins from MCL-1. MCL1 is a very labile protein with a half-life protein around 30 min [13,24] stabilized in many cell types by occupancy of its BH3 binding grove by endogenous proteins or S63845 [21–23,34,35]. This also occurs in bCAFs as S63845 induces an increase in MCL-1 protein expression level and further studies are required to identifiy bCAF expressed proteins involved in MCL-1 turn-over. Stabilization of MCL-1 in S63845 treated bCAFs coincides with its interaction with DRP-1 in a Mdivi-1 sensitive manner. MCL-1 was already described to interact with DRP-1 (in addition to OPA1), for instance in human pluripotent stem cells (hPSCs) and in hPSC-derived cardiomyocytes [36,37] [29,30]. The most likely explanation for the S63845 effect in bCAFs is that the treatment allows grove independent interaction of MCL-1 with DRP-1 at mitochondria (where MCL-1 was found to accumulate) leading to organelle fission and subsequent loss of myofibroblastic features. In support to this hypothesis, we found that MCL-1 knock down (KD) in bCAFs has sensitized them to BCL-xL inhibitor (supplementary Figure 2A) but did not phenocopy the effects of S63845 on mitochondrial fragmentation and stress fiber loss by itself and lost the response to S63845 (supplementary Figure 2B). Moreover, MCL-1 KD bCAFs cells showed no alteration in invasiveness but lost their response to S63845 (incidentally confirming that the effects of S63845 are on target, supplementary Figure 2C). Of note, MCL-1-silenced embryonic stem cells exhibited fused mitochondria[29] arguing for an active, possibly grove independent role for MCL-1 in mitochondrial fission in these cells. Consistently, the effects of S63845 do not necessitate BAX-BAK expression: MCL-1 targeting diminished stress fibers in BAX-BAK KD bCAFs (supplementary Figure 2B), note that these cells are resistant to S63845 combined with A1331852 (supplementary Figure 2A). Regardless of the underlying mechanisms involved in S63845-induced mitochondrial fragmentation, our study unravels a link, in bCAFs, between mitochondrial fusion/fission dynamics, stress fibers and the expression of a set of myofibroblastic genes. These are known to be regulated by transcription factors such as SRF-MRTF and YAP-TEAD whose mechano-dependent activity is itself dependent on an active actomyosin network [11]. We propose that S63845 treatment interferes with this positive feedback loop, as it prevented the nuclear localization of YAP in the same time as it prevented stress fiber formation. Numerous studies have established an influence of mitochondrial function on cytoskeleton organization and/or mechano-dependent transcription factors [38,39]. It should be noted that we observed, during S63845 treatment, changes in myofibroblastic features before we could detect any change in mitochondrial mass or oxidative phosphorylation. We thus propose that the biological effect we describe relies on a more direct link between the mitochondrial fusion-fission machinery and the cytoskeleton.

CAFs are an heterogenous component of the cellular microenvironment as underlined by numerous studies identifying phenotypically distinct CAFs subsets in various cancers [3,19,40–43] There seems to be some consensus regarding the existence of perivascular CAFS and of ECM rich CAFs. The latter population is enriched by our isolation protocol [13], and heterogenous in itself. Breast ECM-CAF encompass, in molecular subtype dependent and patient specific proportions, CAF subsets endowed with an inflammatory phenotype (iCAFs) and other myofibroblastic CAFs (myCAFs). A similar dichotomy was described in pancreatic ductal adenocarcinoma models and clinical samples, where IL1-driven iCAFs and TGFb driven myCAFs were described [44]. The latter study unravels an activation process with a transition from normal fibroblasts to a single CAF phenotype, later diverging to either iCAF or my CAf. myCAFs are of particular relevance as they exert an immunosuppressive effect and are associated with primary resistance with immunotherapy. Our study indicates a strong dependency on MCL-1 regulated mitochondrial fusion/fission dynamics of the myCAF phenotype. Further studies are required to determine at which step(s) of the CAF activation process MCL-1 is involved. In particular, understanding whether MCl-1 expression and interactome are differently regulated in distinct subsets during activation is of particular interest. Maybe one of the most exciting implication from our study is that it shows that activation may be reversible, as myofibroblastic features can be pharmacologically reversed using a small molecule targeting MCL-1. This pleads for the use of such a compound for the treatments of cancers with stromal evolutions into myofibroblasts (with advantages and limitations discussed above) but also give indications about the influence of current treatments might have on the tumor stroma. It is relevant here to note that MCL-1 mRNA and protein expression levels are inhibited by anthracyclines, which are part of the chemotherapeutic cocktail used for breast cancer therapy. Analyzing the clinical effects of chemotherapy on CAF expressed MCL-1 may thus help predict tumor response.

## ACKNOWLEDGMENTS

We thank members of the “Stress adaptation and tumor escape” laboratory for their support. We thank Drs V. Maguer-Satta, C. Blanquart and V. Sauzeau for fruitful discussions and their suggestions. We thank P. Gomez, C. Seiller and Gavard-Bidère team members for providing reagents and technical advice. We benefited from technical support from the cytometry (CytoCell). We acknowledge the MicroPICell facility, SFR-Santé, INSERM, CNRS, UNIV Nantes, CHU Nantes, Nantes, France, member of the national infrastructure France-BioImaging supported by the French National Research Agency (ANR-10-INBS-04), special thanks to P. Hulin and S. Nedellec for their advice. We are most grateful to the Genomics and Bioinformatics Core Facility of Nantes (GenoBiRD, Biogenouest, IFB) for its technical support. T.L Bonneaud was supported by a *fellowship* from Ligue contre le cancer (CD44 and CD36), A. Basseville benefited from the financial support of European Commission with Marie Sklodowska-Curie Action research grant and L. Nocquet was a recipient of French ministry of Higher Education, Research and Innovation. This work was supported by INCa and DGOS (SIRIC ILIAD, INCa-DGOS Inserm-12558), Ligue contre le cancer (CD44, CD53 and CD22) and Run Odyssea to ICO-Gauducheau.

