## Supplementary Material for "Targeting of MCL-1 in breast cancer associated fibroblasts reverses their myofibroblastic phenotype and pro-invasive properties"

**Supplementary Figure 1.** **A** Relative Oxygen Consumption Rate (OCR) measured by seahorse in CAFs treated with S63845 (500nM) compared to the control (untreated). Data are means ± SEM from three independent experiments. P value was determined by student t-test. ns: not significant. **B** TOM-20 protein expression levels in primary culture of CAFs treated with S63845 500nM for 18 hours were evaluated using western blots analysis. Actin protein expression was used as loading control. **C** MCL-1 protein expression in CAFs treated by S63845 (500nM) or A1331852 (100nM). **D** Mitochondrial colocalization of TOM20 (red) and MCL-1 (green) in CAFs treated or not by S63845 or A1331852.

**Supplementary Figure 2.** **A-B** CAFs sg control or silenced for MCL-1 (sg MCL-1) or BAX and BAK (sg BAX-BAK) were treated or not with S63845 (500nM), A1331852 (100nM) or S63845 (500nM) + A1331852 (100nM) for 48 hours in DMEM containing 1% FBS. **A** Top: Protein expression level were evaluated using western blots, bottom: apoptosis was measured by Annexin-V flow cytometry. Data are means ± SEM from three independent experiments. P value was determined by two-way ANOVA. ****P<0.0001, ns: not significant. **B** Quantification of F-actin staining with FITC-phalloidin and immunofluorescence of TOM-20 (fused or fragmented mitochondria) in CAFs Sg control or sg MCL-1 treated or not with S63845. Randomly generated images set names were randomized for analysis. Around sixty to one hundred cells were analyzed per condition. Counting events were done manually through NIS-Elements software (Nikon software). Fluorescence intensity and positivity staining were determined by positive and negative control comparison, dependent on the experiment. **C** 3D spheroid of CAFs sg control or sg MCL1 invasion into collagen in presence or not of S63845 (500nM) was monitored up to 48h using microscopy time-lapse. Representative images of tumor cell invasion are shown.

Bonneaud et al, supplementary Fig 1

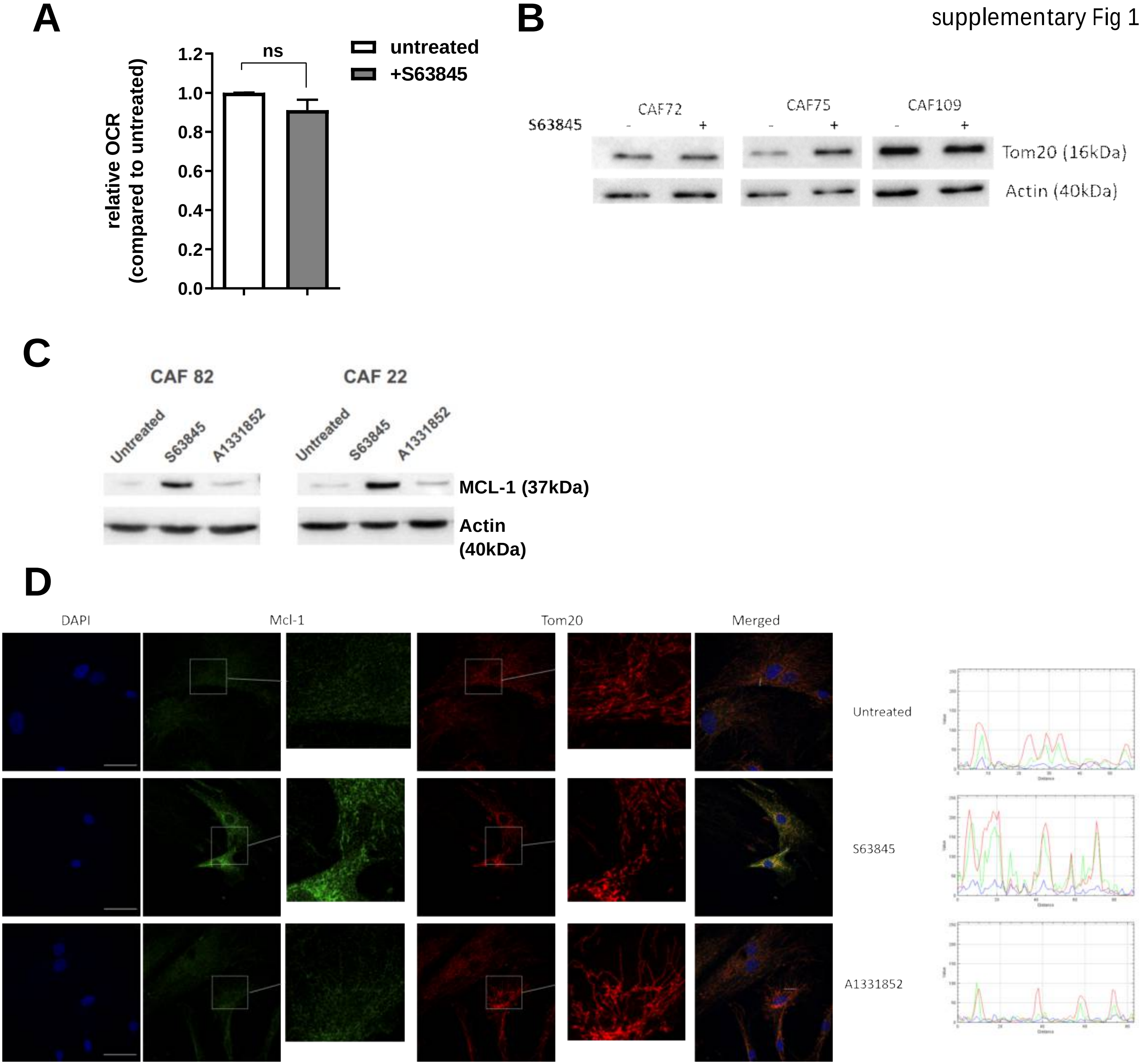

supplementary figure 2

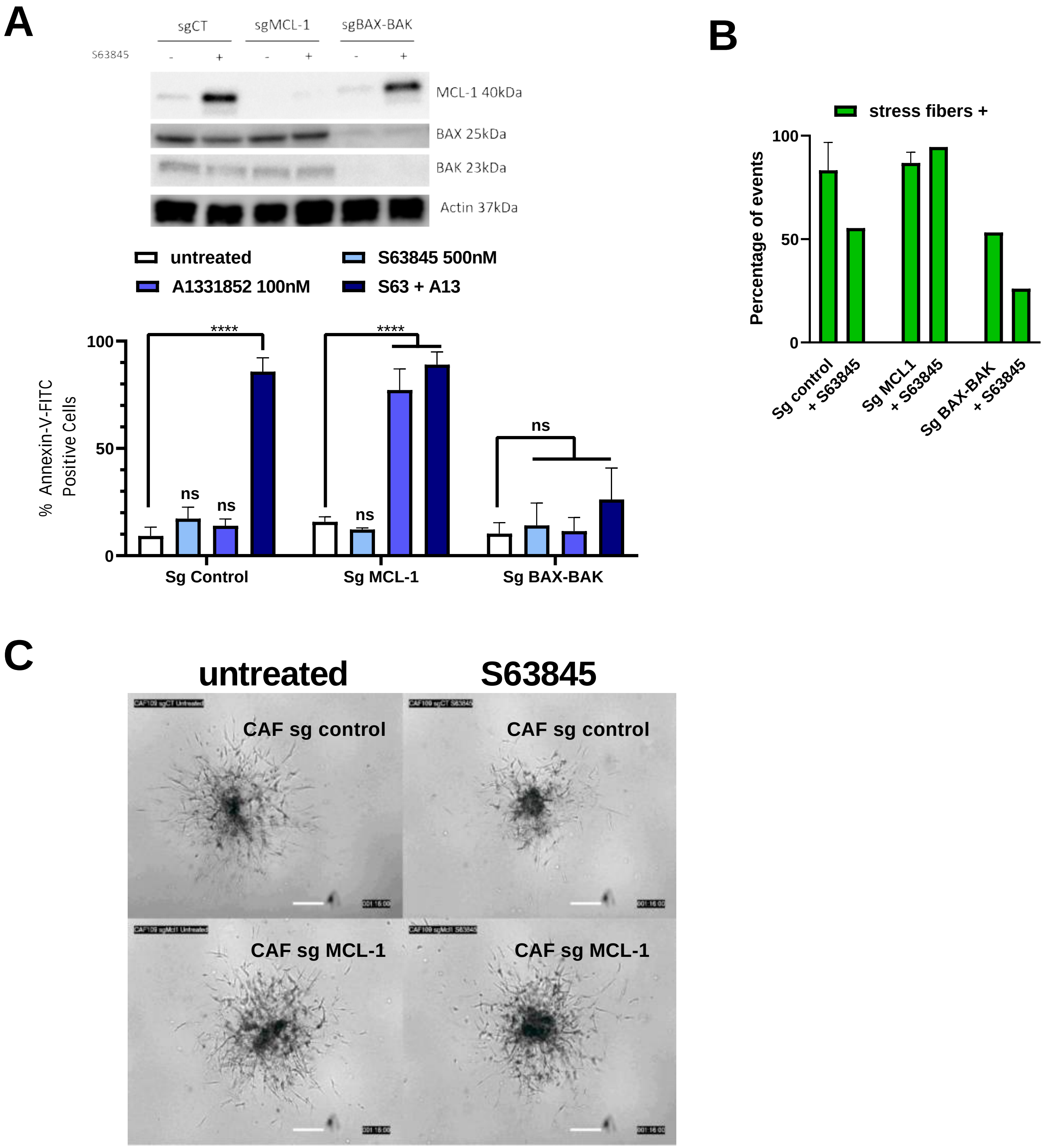
